# Nanoscopic Structural Fluctuations of Disassembling Microtubules Revealed by Label-Free Super-Resolution Microscopy

**DOI:** 10.1101/2020.10.21.348904

**Authors:** Milan Vala, Łukasz Bujak, Antonio García Marín, Kristýna Holanová, Verena Henrichs, Marcus Braun, Zdeněk Lánský, Marek Piliarik

**Affiliations:** Institute of Photonics and Electronics of the Czech Academy of Sciences, Chaberská 1014/57, 182 51 Prague, Czech Republic; Institute of Biotechnology of the Czech Academy of Sciences, BIOCEV, Průmyslová 595, 252 50 Vestec, Czech Republic

**Keywords:** microtubule, scattering anisotropy, dynamic instability, iSCAT, super-resolution microscopy

## Abstract

Microtubules are cytoskeletal polymers of tubulin dimers assembled into protofilaments that constitute nanotubes undergoing periods of assembly and disassembly. Static electron micrographs suggest a structural transition of straight protofilaments into curved ones occurring at the tips of disassembling microtubules. However, these structural transitions have never been observed and the process of microtubule disassembly thus remains unclear. Here, a label-free optical microscopy capable of selective imaging of the transient structural changes of protofilaments at the tip of a disassembling microtubule is introduced. Upon induced disassembly, the transition of ordered protofilaments into a disordered conformation is resolved at the tip of the microtubule. Imaging the unbinding of individual tubulin oligomers from the microtubule tip reveals transient pauses and relapses in the disassembly, concurrent with enrichment of ordered protofilament segments at the microtubule tip. These findings show that microtubule disassembly is a discrete process and suggest a mechanism of switching from the disassembly to the assembly phase.

## 1. Introduction

Experimental characterization of the dynamics of a biological system provides essential knowledge necessary to understand its function. Currently, snapshots of biomolecular assemblies can be obtained at an atomic resolution by cryogenic electron microscopy (cryo-EM).^[1,2]^ However, these conventional structure-resolving methods do not provide any information on the dynamics of the assemblies. Optical microscopy remains the method of choice for deciphering fast dynamic processes in biological systems, and, in combination with sparsely labeled structures, enables visualization of the positions of single molecules with nanometer resolution.^[3,4]^ Nonetheless, detailed macromolecular arrangements and structural fluctuations remain unresolvable by direct optical means, hindering our understanding of the dynamics of biological systems.

Microtubules (MTs) are highly dynamic biopolymers, constituting one of the key components of the eukaryotic cytoskeleton. An MT typically consists of 13 chains (protofilaments) of head-to-tail-connected tubulin dimers that laterally interact to form a hollow cylinder approximately 25 nm in diameter.^[5,6]^ MTs stochastically undergo sudden transitions between phases of slow assembly (polymerization) and rapid disassembly (de-polymerization).^[7–9]^ This dynamic instability is vital for the multiple roles of MTs in cells, including cell division, motility or intracellular transport,^[10]^ and is regulated by additional factors during different phases of the cell cycle^[11–13]^ or by drugs, including the anticancer compound Taxol.^[14]^ Although vital, the process of MT disassembly remains elusive, mainly due to the lack of information on the dynamics of the underlying structural changes at the MT tip. MT disassembly is attributed to the release of strain energy accumulated in the microtubule lattice, where the intrinsically curved GDP tubulin dimers are held in a straightened, and thus strained, conformation.^[15–18]^ This strain release at the MT tip is believed to result in a conformational change in tubulin dimers relaxing into their curved conformation, which results in a formation of curved protofilament structures formed by short tubulin oligomers at the MT tip^[19]^, and consequent unbinding of the oligomers from the MT. In a domino-like effect, the successive unbinding of tubulin oligomers at the MT tip results in the MT disassembly. Although the force generated by the disassembling tip was recently measured^[20]^, the dynamics of the process is still not known.

The vast majority of studies of MT dynamics are based on fluorescence microscopy, capable of the quantification of MT (dis)assembly rates and monitoring the interaction with MT-associated proteins.^[13,21,22]^ MTs are known to be optically anisotropic, allowing for their label-free observation with polarization microscopes.^[23–25]^ A highly sensitive method for imaging weakly scattering objects, including single MTs, is interferometric detection of scattering (iSCAT).^[3,26,27]^ iSCAT is based on the common-path interference of the scattered light with the reference wave, allowing for imaging and tracking of nanoparticles^[3]^ as small as 2 nm^[28]^ and the detection of single unlabelled proteins.^[26,29]^ iSCAT was recently used to track the displacement of single unlabelled MTs with sub-nanometer precision^[30]^ and to observe the binding of nanoparticle-labeled tubulin dimers to growing MT ends.^[31]^ However, the conformation of single proteins or short protein oligomers, such as tubulin oligomers at the disassembling MT tip, has never been optically resolved by direct and label-free means.

Here, we introduce a new label-free light microscopy technique capable of distinguishing macromolecular structures based on their immediate conformation and explore the rapid dynamics of tubulin oligomers during MT disassembly by directly visualizing the dynamics of the wave of tubulin conformational changes at the disassembling MT tip. The experimental approach is based on a parallelized polarization-sensitive iSCAT microscope and using the polarization-dependent contrast to estimate the immediate fraction of curved tubulin dimers and the length of the transiently disordered region at the disassembling MT tip. We show that the disassembly velocity fluctuates in time, exposing millisecond periods of disassembly pause or relapse. Localization of single events of tubulin oligomer unbinding renders a superresolved image of the MT without the need for any artificial labels and, at the same time, enables the colocalization of the unbinding events with the preceding conformation changes, all taking place within a diffraction-limited range from the microtubule tip.

## 2. Results

### 2.1. Polarization-sensitive iSCAT microscopy

The experimental setup is based on a conventional wide-field iSCAT microscope^[26]^ that was extended to examine the same field of view at two orthogonal polarizations of light in a single camera frame (**Figure 1a**). A similar arrangement was previously used to detect the orientation of metallic nanorods using both iSCAT^[32,33]^, and dark-field^[34]^ microscopy; however, here we implemented a method that allows single protein sensitivity^[26]^. We recorded a series of images of single depolymerizing MTs, oriented parallel or close to parallel (±10 degrees) to one of the detected polarizations, with a temporal resolution of 380 μs. In the case of an anisotropic scatterer such as an MT (Figure 1c), the scattered field *E_s_* splits into the detected polarizations 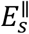 and 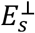, resulting in different iSCAT contrasts in the two respective polarization channels (Figure 1b and Supporting Information).

**Figure 1.**
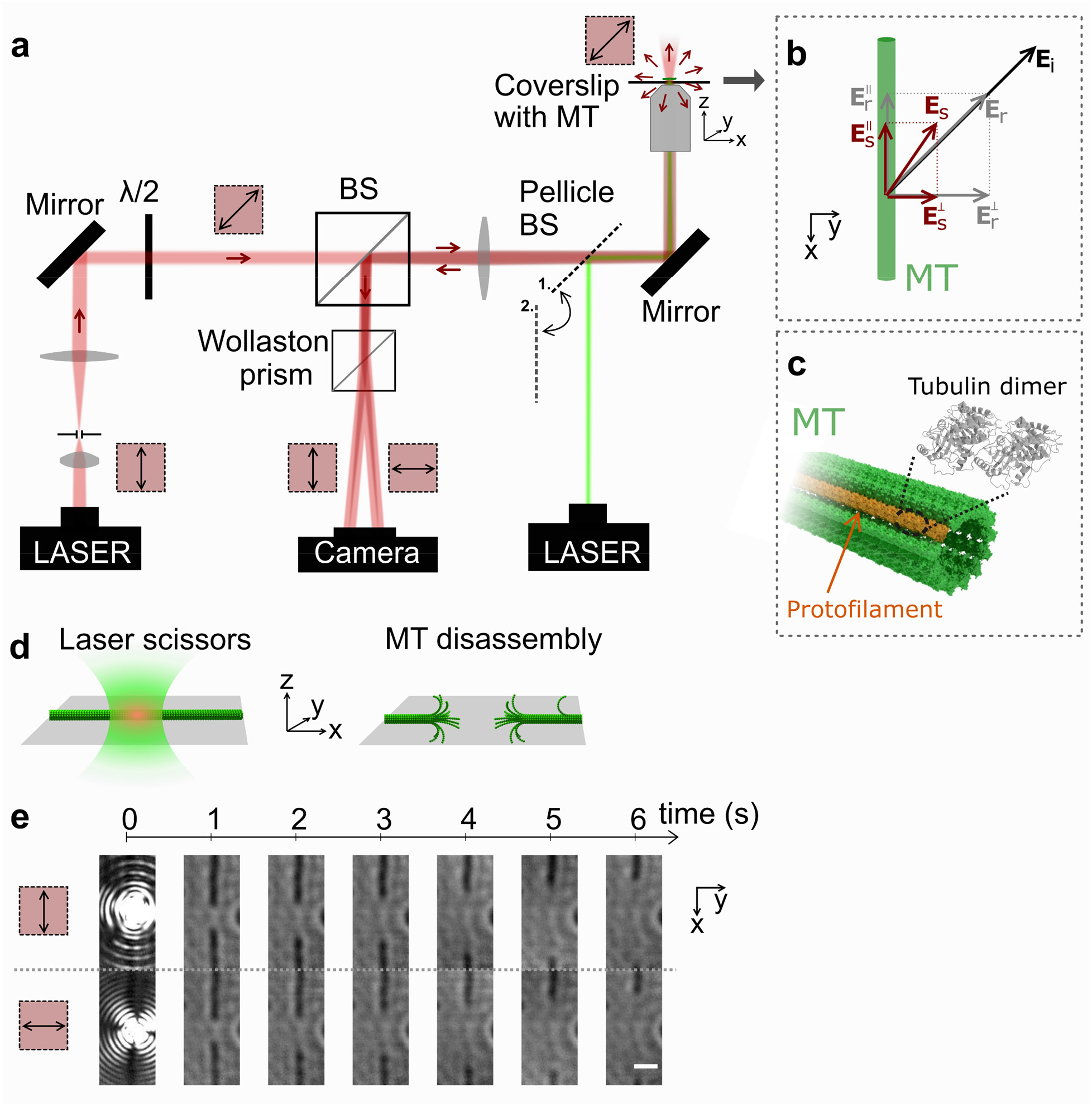
Monitoring the disassembling MT with the polarization-sensitive iSCAT microscope. (**a**) The layout of the modified polarization-sensitive iSCAT microscope for monitoring of anisotropy fluctuations. (**b**) Scheme of the amplitudes of the scattered and reflected signals and their decomposition into waves linearly polarized parallel and perpendicular to the MT. (**c**) Illustration of the MT structure with the tubulin dimer subunits arranged in the cylindrical lattice. (**d**) Illustration of the MT disassembly initiated by the laser scissors triggered at position 1 of the pellicle BS. **(e)** Time-lapse iSCAT images of a single MT acquired at two orthogonal polarizations after the laser-scissors-induced disassembly (t=0). The scale bar corresponds to 1 μm.

We polymerized GDP MTs, which were stabilized at their tips by caps formed by tubulin bound to a non-hydrolyzable GTP analog guanosine-5’-[(α,β)-methyleno]triphosphate (GMPCPP), and attached them specifically to the coverslip surface. A second laser beam (*λ* = 532 nm, Figure 1a) was focused by the microscope objective on a non-stabilized GDP segment of the MT (Figure 1d) resulting in the scission of the MT and formation of two freshly exposed tips (time t = 0 s) that immediately started to disassemble. A series of iSCAT images capturing the laser scissors and subsequent disassembly were recorded for up to 16 s in both recorded polarizations as illustrated in Figures 1d,e. A sample video of the process recorded at 2900 frames per second is available in Movie S1.

### 2.2. Resolving changes in nanoscopic structures at the disassembling MT tip

The two orthogonal projections of the detected signal represented by the two contrasts c^‖^ and c^⊥^ characterize the amplitude and polarization of the scattered light and directly define the real part of the Jones polarization vector (c^‖^, c^⊥^).^[35]^ This representation allowed us to filter out an image of known scattering anisotropy contributing to the overall signal using a simple linear projection. In this procedure, first, we spatially overlapped the images acquired at the two respective polarizations (**Figure 2a**) with a precision of approximately 5 nm. Next, we used the aligned pairs of images to calculate the mean contrast of each disassembling MT tip, which is presented in Figure 2b in the form of a kymograph of the cross-section along the length of the MT. The profile of the mean contrast (c^‖^ + c^⊥^)/2, i.e., an unpolarized iSCAT contrast, can be associated with the distribution of the tubulin mass along the MT.^[29]^ To highlight the contribution of the anisotropic scattering, we calculated the differential contrast (c^‖^ − c^⊥^). The kymograph constructed from such a differential signal, in which the contribution of the isotropic scattering cancels out, is shown in Figure 2c. In our experiment, the anisotropic scattering of the stable MT was well-characterized by the scattered polarization vector 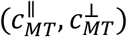 obtained at the beginning of each experiment. Consequently, we subtracted the contribution of the anisotropic scattering of the stable MT from the resulting image as 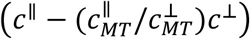. An increase in the isotropic signal at the tips of the disassembling MT was observed in the resulting kymograph (Figure 2d). Both the anisotropic and isotropic contributions are highlighted in a composite, false-color kymograph in Figure 2e.

**Figure 2.**
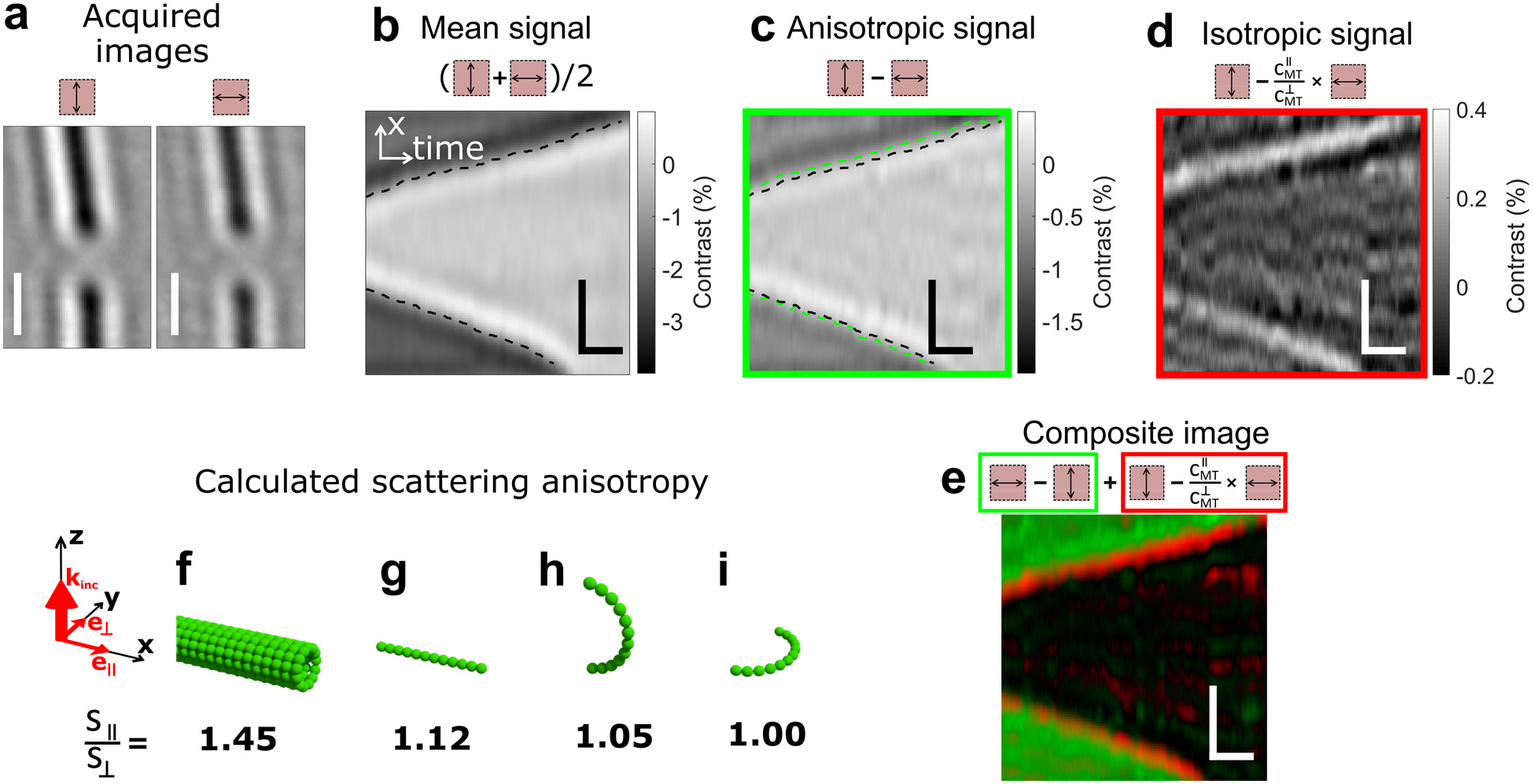
Scattering anisotropy of a disassembling MT end. (**a**) Images of MT acquired with light polarized parallel (along the **x**-axis) and perpendicular to (along the **y**-axis) the MT. (**b**) Kymograph of the iSCAT contrast (mean of the contrasts measured in both polarizations) of the disassembling MT. (**c**) Kymograph of the anisotropic and (**d**) isotropic contribution to the overall MT contrast. (**e**) The composite kymograph of isotropic (red) and anisotropic (green) scattering combined as a false-color image. Scale bars in (**a-e**) correspond to 1 μm (vertical) and 1 s (horizontal). Illustration of different MT substructures and their numerically calculated ratios of scattering anisotropy 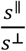: (**f**) complete MT shaft, (**g**) straight protofilament, (**h**) protofilament curved in axial, and (**i**) in lateral microscope plane.

The comparison of the kymographs in Figure 2 shows that the gap between the microtubule tips in the kymograph of the anisotropic signal (green dashed line in Figure 2c) is slightly wider than the gap in the kymograph of the mean signal (black dashed line in Figure 2b,c). This observation indicates that the decrease in the scattering anisotropy of the MT precedes the loss of mass at the very tip of the disassembling MT. We attribute the local increase in the isotropic signal to the presence of tubulin dimers that are still attached to the tip of the MT, but whose lower optical anisotropy is caused by a loss of their ordered arrangement in the MT lattice. We support this assumption with a set of finite-difference time-domain (FDTD) simulations of the scattering anisotropy of the whole MT shaft and several possible conformations of protofilaments, provided in detail in the Supporting Information. Modeled geometries with calculated scattering anisotropies are illustrated in Figure 2f-i, which imply that the conformation change of a single protofilament from straight to curved results in a decrease in its scattering anisotropy. Combined, these results demonstrate that our polarization-sensitive iSCAT method enables the resolution of nanoscopic structural changes at the tip of a disassembling MT and discern the image of the structured MT shaft from the image of the unstructured region at the MT tip.

### 2.3. Estimation of the length of the unstructured region at the disassembling MT tip

Next, we quantified the axial displacement between the overall image of the disassembling MT and the corresponding anisotropic contribution, which we can interpret as the mean length *Δx_c_* of the unstructured region at the end of the MT. The averaged spatial profiles of the mean iSCAT signal and anisotropic signal over time for each of the recorded disassembling MT tips is shown in **Figure 3d**. The vast majority of the measured disassembling MTs feature a distinct offset between the mean and differential contrast. We fitted the profiles of mean and differential contrast with a sigmoidal function and calculated the displacement of the sigmoid centers of the observed disassembling MT ends, yielding a mean displacement of *Δx_c_* = (78 ± 39) nm (Figure 3g).

**Figure 3.**
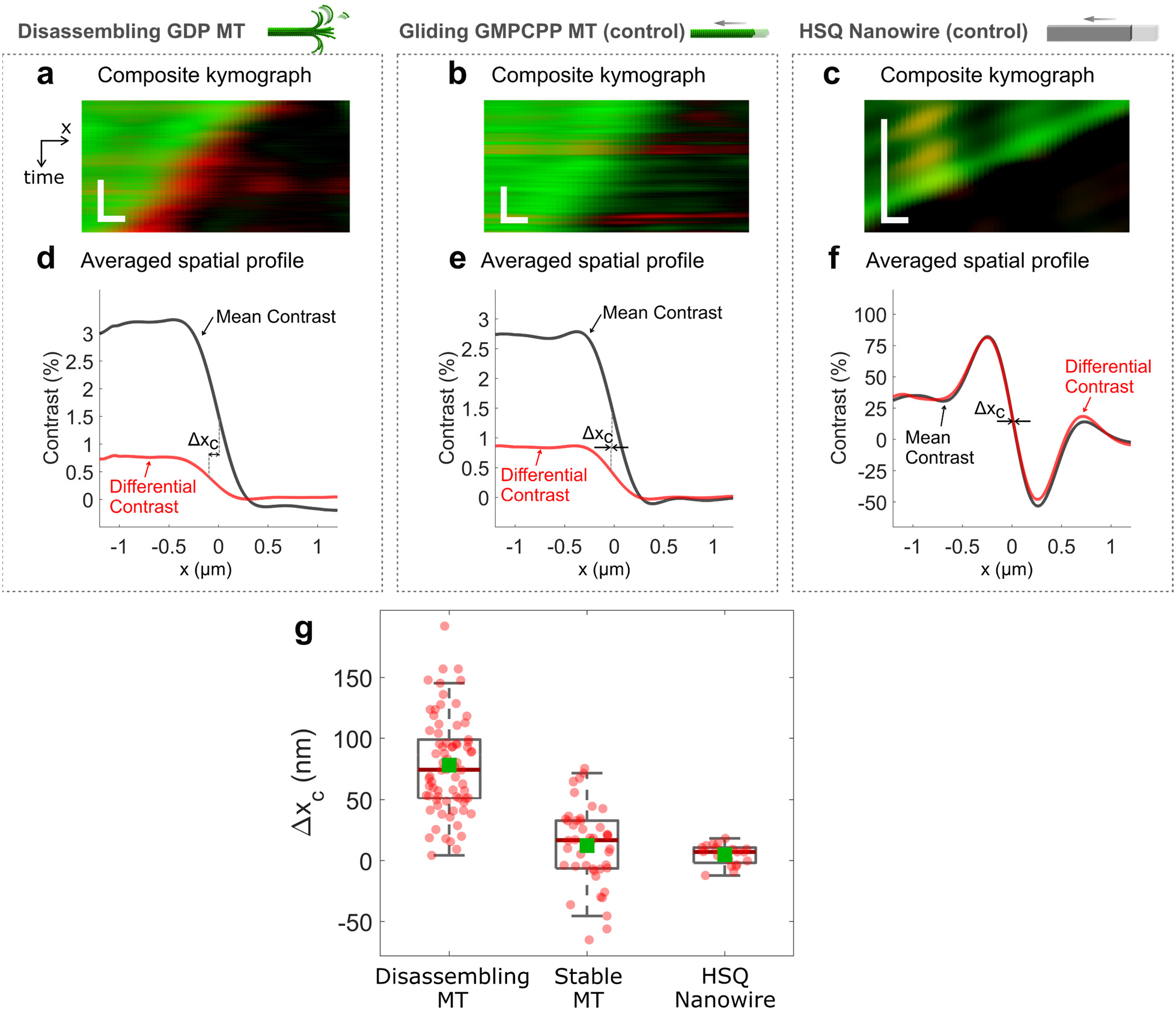
Displacement of the differential and mean scattering contrasts of a disassembling MT and two controls. Composite kymographs of the anisotropic (green) and isotropic contribution (red) to the (**a**) disassembling MT contrast, (**b**) gliding double-stabilized MT, and (**c**) dielectric nanowire. Scale bars correspond to 1 s (vertical) and 200 nm (horizontal). (**d**) The profile of the mean contrast (black line) and anisotropic contrast (red line) averaged from disassembling MTs, (**e**) double-stabilized MTs, and (**f**) dielectric nanowires. (**g**) Box plot summarizing the displacement *x_c_* between the centers of the edges of the mean and anisotropy profiles for disassembling MT tips (N=74) and both controls (stable MT, N=44; nanowire, N=20). Red lines and green squares correspond to median and mean values, respectively.

We performed two control experiments in which no or negligible structural change was expected. In the first control, a double-stabilized MT was attached on the surface via immobilized kinesin-1 molecular motors. Instead of triggering the MT disassembly, ATP was introduced into the channel to trigger the kinesin-propelled gliding of the MT along the surface^[36]^. Imaging the stable (non-disassembling) tip of the gliding MT resulted in a set of data visually similar to the retreating tip of a disassembling MT (Movie S2). Very small changes of the contrast anisotropy were observed near the gliding MT tip (Figure 3b) and the displacement parameter *Δx_c_* of (12 ± 33) nm (Figure 3g) falls within the experimental error, which is also similar to the experimental error of the *Δx_c_* of the disassembling MT. For the second control, we used the nanofabricated dielectric nanowires (see Methods and Figure S2), which were moved with the substrate at a constant velocity and assessed the precision of the displacement estimation (Figure 3c and 3f). The statistics of the fitted displacement *Δx_c_* = (5 ±8) nm shown in Figure 3g falls within the experimental error of the spatial alignment of the images taken at different polarizations.

A detailed discussion and interpretation of the displacement *Δx_c_* is provided in the Supporting Information; however, in the simplest model, we directly associate the *Δx_c_* with the mean length of the loose protofilaments in the curved conformation. The quantitative result obtained for the disassembling MT of approximately 10 tubulin dimers closely matches the mean length of curved protofilaments observed in previous electron-microscopy-based studies^[37]^.

### 2.4. Microtubule disassembly rate correlates with the amount of curved protofilaments

We analyzed the changes in the size of the unstructured region of the MT tip and investigated its relation to the MT disassembly rate. We tracked the immediate position of the disassembling MT end with the localization precision *σ_x_* = 2.6 nm and temporal resolution of 380 μs. Two examples of MT end trajectories are shown in **Figure 4a**. The high-resolution trajectory revealed a fluctuating character of the disassembly when compared with a much smoother trajectory obtained from the gliding MT control presented in Figure 4b. We calculated the instantaneous velocity of the receding MT tip (instantaneous disassembly rate) with an accuracy better than *σ_v_* = 10 nm s^−1^ (see Figure S6 for exemplar time series of velocities and histograms of velocities for all 34 pairs of MT ends). We note that (in the absence of tubulin dimers in solution) the disassembly process pauses several times per second (Figure 4a) and the velocity of the receding tip can occasionally reach negative values (Figure 4c). Statistics of receding tip velocity summarized from the two disassembling ends of a single MT and the full set of disassembling MTs (sorted into two groups of slower and faster ends from each pair of MT tips) are compared to the velocity histogram of a gliding MT in Figure 4c. The polarization-sensitive iSCAT microscopy enables simultaneous measurement of the tip velocity and the magnitude of the isotropic component of scattering at the MT tip, as illustrated in an example in Figure 4d-f. Interestingly, the MT tip velocity correlates with the magnitude of the isotropic signal in this example, as well as when cross-correlated and averaged through all the data analyzed (Figure 4g). The relation between the contrast of the unstructured region and the MT tip velocity is visualized in the 2D histogram in Figure 4h. To highlight the difference between the limit cases of progressing or stalled disassembly, the magnitudes of the isotropic signals were sorted into two groups based on the corresponding instantaneous velocities being positive or non-positive, respectively (Figure 4i). Combined, these data provide direct evidence showing that the rate of the MT tip disassembly correlates with the amount of isotropically scattering, likely curved, protofilament structures at the MT tip. The fluctuating character of the MT disassembly resolvable at high acquisition rates is presumably related to the discrete mechanism of the MT disassembly, in which segments of protofilaments of various lengths unbind from the disassembling tip.

**Figure 4.**
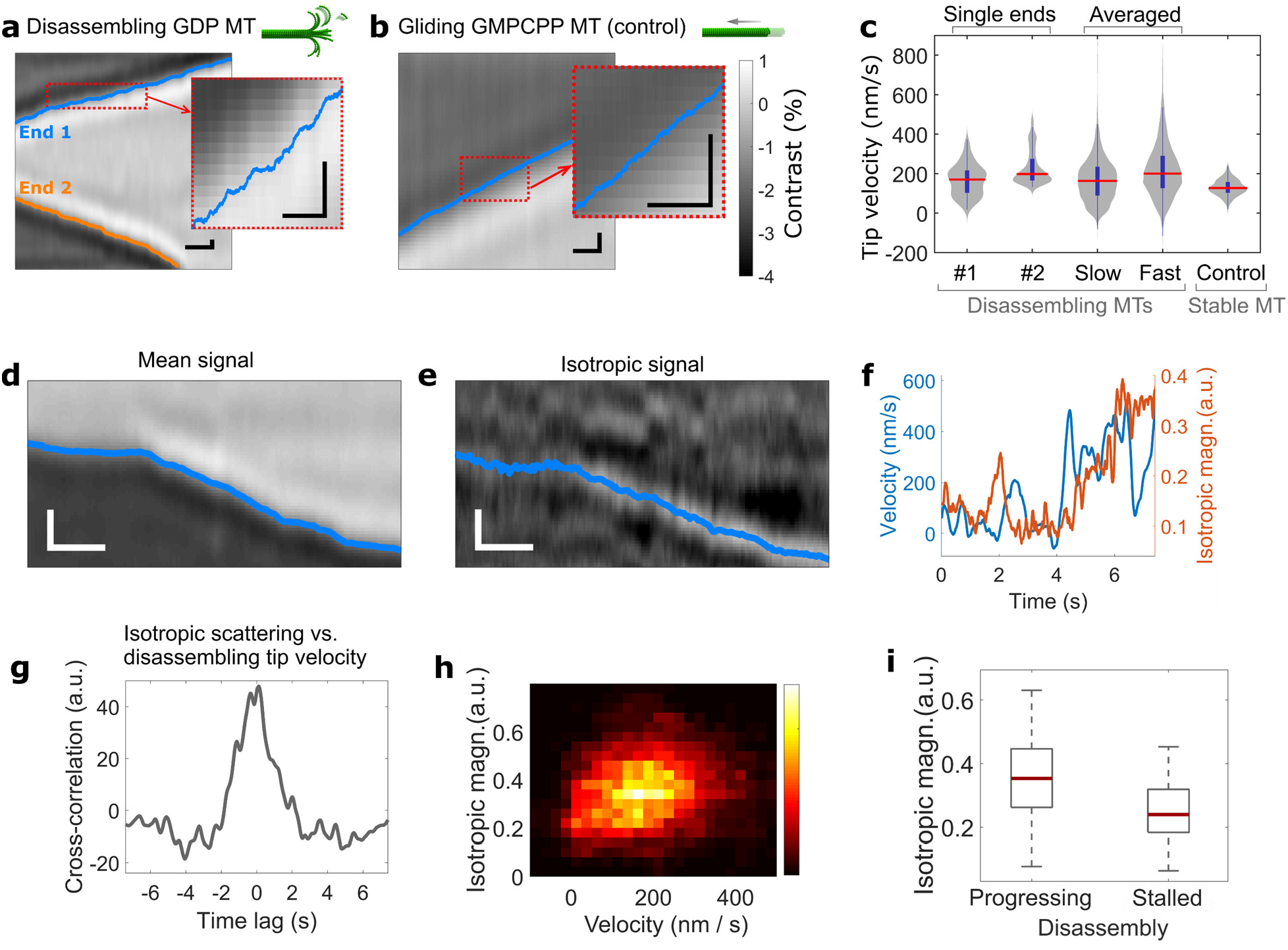
Fine structure of the disassembly rate correlates with the molecular disorder at the MT tip. (**a**) Kymograph of a mean iSCAT signal with localized positions of the MT ends. The red dashed line highlights the zoomed region. (**b**) Identically processed kymograph of the gliding double-stabilized MT. Scale bars correspond to 1 s (horizontal) and 200 nm (vertical). (**c**) Histograms of the velocities of the tracked MT tips. From left to right, two individual ends of the disassembling MT from (**a**), accumulated velocities from slower and faster ends from each of 34 pairs of measured disassembling MT ends, and velocity of individual gliding MT end from (**b**). Example of a kymograph of (**d**) mean and (**e**) isotropic signal of another disassembling MT end, and (**f**) corresponding plot of a disassembling tip velocity and the magnitude of an isotropic signal. (**g**) Measured cross-correlation between the immediate end velocity and magnitude of the isotropic scattering. (**h**) 2D histogram of isotropic signal magnitude and end velocity. (**i**) Comparison of isotropic signal magnitudes measured at periods of progressing (*v_disassembly_*>0) and stalled (*v_disasembly_*<0) disassembly. Data in (**g-i**) were obtained from n = 68 disassembling MT ends.

### 2.5. Label-free super-resolution imaging of the disassembling MT

The fluctuating character of the velocity of the receding MT tip is also visible in the incremental kymograph in **Figure 5a** compiled from series of incremental images (two examples in Figure 5b,c) made by subtraction of the iSCAT images taken at times *t* and *t* + *Δt* (*Δt* = 19 ms in Figure 5). Clearly, the contrast fluctuation in the incremental kymograph correlates closely with the instantaneous disassembly rate. The minimum detectable contrast above the background noise (3σ condition) had an amplitude of 0.015%, corresponding to the unbinding of approximately 3 tubulin dimers from the MT tip (see Methods). A histogram of contrasts of all the peaks detected in the incremental images is shown in Figure 5g, revealing that the mean size of the detected unbound protofilament segment is approximately 8 tubulin dimers. The sum of all the diffraction-limited peaks reaching up to 60% of the whole MT signal is shown in Figure 5d, indicating the majority of the tubulin oligomer unbinding events were detected.

**Figure 5.**
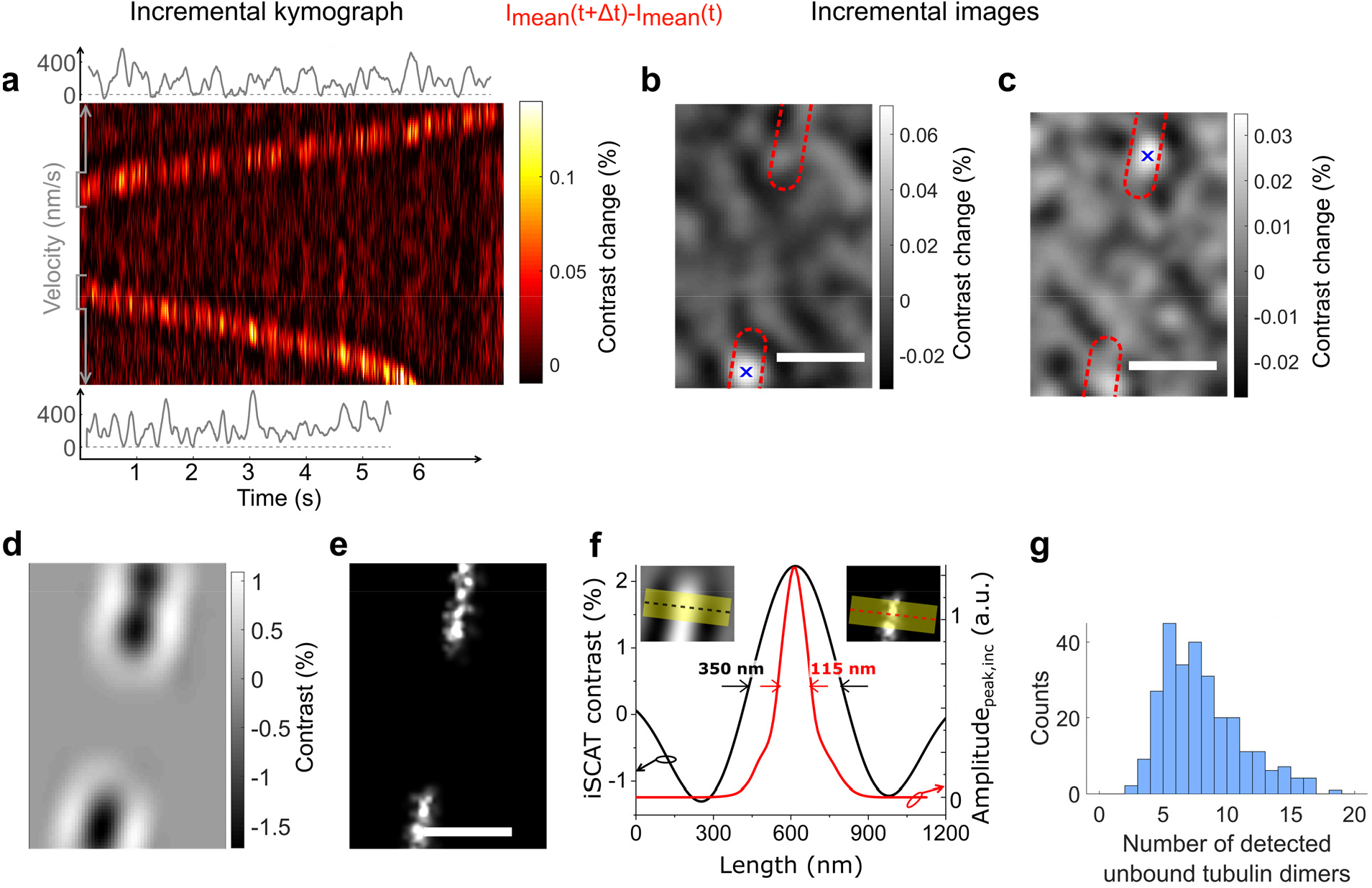
Label-free super-resolution reconstruction of the disassembling MT image. (**a**) Incremental kymograph of the disassembling MT constructed for temporal increment *Δt* = 19 ms. Corresponding velocities of the disassembling ends are shown above and below the kymograph. Examples of incremental images with the detected local maxima caused by the loss of mass from (**b**) lower and (**c**) upper MT tip. The blue cross highlights the peak center localized using a Gaussian fit. (**d**) Image of the segment of the MT disassembled within 4.8 s reconstructed from detected peaks convolved with PSF of the iSCAT microscope, and (**e**) the corresponding super-resolved image reconstructed from detected peaks. (**f**) Cross-sections of the iSCAT and super-resolved MT images. (**g**) Histogram of the contrasts of the detected incremental peaks expressed in units of tubulin dimers unbound from the MT end.

We localized the detected peaks in the incremental images with a precision of 20 nm or better (depending on the contrast^[38]^) and reconstructed the super-resolution image of the disassembled part of the MT shown in Figure 5e. The cross-section profile (Figure 5f) of the incremental super-resolved image shows FWHM_*SR,inc*_ = 115 nm, which is in approximate agreement with the sum of the diameter of the microtubule (25 nm), the displacement of frayed protofilaments (approximately 30 nm on either side of the MT based on the published cryo-EM images^[37]^), and the localization precision.

### 2.6. Detection of conformation changes during the MT disassembly

We overlapped a series of images of disassembling MTs based on the fitted position of the MT tip and accumulated images of the MT; their separation into the isotropic and anisotropic contributions and the corresponding sum of incremental images taken from the respective data are shown in **Figure 6a-f.** Strikingly the bright spot in Figure 6d indicates the position where the contrast is removed from the anisotropic image of the MT (Figure 6c) and transferred to the (isotropic) image of unstructured segments (Figure 6e). This position does not necessarily overlap with the mean displacement of the end of the structured segment of the MT explored in Figure 3 as the transition from structured to unstructured MT might have an arbitrary gradient character (see the linearized model in the Supporting Information). A specific combined pattern of a dip and a peak closely matching the subtraction of the images in Figure 6b and Figure 6d is shown in Figure 6f, indicating the MT mass transfer from the anisotropic conformation to the isotropic conformation and consecutive disassembly of the tubulin mass while it is in the isotropic (kinked) conformation. Furthermore, conformation changes such as MT protofilament curving are expected to be associated with a small momentary displacement of molecular mass along the MT axis, also resulting in a pattern such as that shown in Figure 6f. Therefore, we searched for this distinct pattern in the individual incremental images, and, to avoid any biased selection in noise fluctuations, we performed the pattern detection in the polarization parallel to the MT and averaged only the corresponding images in the perpendicular polarization in Figure 6g (see Methods for more details). For reference, we performed the same pattern detection and analysis with the gliding MT data in Fig. 6h, which indicated no signature of mass displacement. The pattern of conformation changes in Fig. 6g features a strikingly strong, yet sub-shot-noise, contrast of ±4×10^−5^. This contrast reached 10% of the mean contrast of single dissociating tubulin oligomers analyzed in Figure 5. Based on this contrast, we can deduce the mean displacement distance by assuming that the mass displaced corresponds to the average 8-tubulin-dimers long protofilament segment. Using the width of the experimental point-spread-function, we deduced a 35-nm displacement within only a 3.5 ms time interval (10 frames binned for each incremental image) appearing at the mean rate of approximately 80 times per second (see Methods). This observation clarifies the dynamics of protofilament conformational changes with previously unseen details. Our data suggest that the conformation change is not a continuous process in which protofilaments gradually peel away from the MT shaft at a rate of one tubulin dimer (8 nm) per 3 ms (corresponding to the 220 nm s^−1^ disassembly rate). Instead, our data suggest that segments of protofilaments, on average 8-tubulin-dimer long, switch from a straight to a curved conformation and unbind from the MT tip.

**Figure 6.**
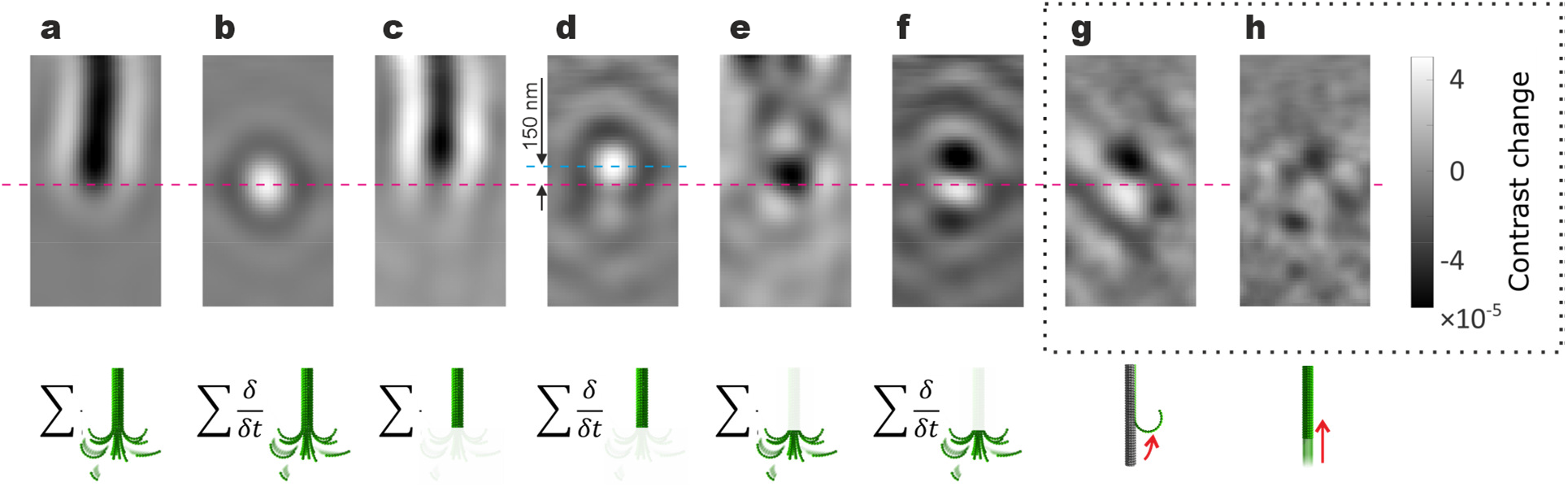
Imaging conformational changes at the tip of the disassembling MT. (**a**) Accumulated iSCAT images and (**b**) incremental images of the tip of disassembling MTs (**c**) Accumulated anisotropic contribution of images of disassembling MTs (structured segment) and (**d**) corresponding incremental image. The blue line denotes the peak position. (**e**) Accumulated isotropic contribution of images of disassembling MTs (unstructured segment) and (**f)** corresponding incremental image. All accumulated images have an arbitrary color scale. (**g**) Mean of mass displacement patterns detected in the incremental images of disassembling MTs and (**h**) gliding MT. The pink dashed line indicates the position of the MT edge.

## 3. Conclusions

We present an ultra-sensitive label-free optical microscopy method capable of capturing rapid changes in the nanoscale geometry of disassembling MT tips. This is achieved by exploring the scattering anisotropy of tubulin oligomers at the disassembling MT tips at submillisecond timescales. Our data provide quantitative information about the immediate cumulative length of MT protofilaments undergoing conformational changes. Our method allowed us to reconstruct a super-resolved image of the disassembling MT in a way similar to stochastic optical reconstruction microscopy (STORM), but without the need for labeling and with high efficiency of the detection of unbinding tubulins reaching up to 60%. This imaging revealed the frayed shape of the disassembling MT tip, suggested before by EM imaging. We were able to discern the delay between protofilament conformational changes and tubulin detaching from the microtubule and to observe long segments of protofilaments, closely matching the mean length of the dissociating tubulin oligomers, coiling momentarily from the MT shaft. These results are in agreement with the current understanding of the geometry of disassembling MTs derived so far solely from static electron micrographs^[37,39]^, supporting the hypothesis that during MT disassembly, segments of protofilaments switch from straight to curved conformation, subsequently detaching from the microtubule tip.

Importantly, we identified fluctuations in the length of the unstructured region at the tip of disassembling MTs and directly showed that faster MT disassembly correlates with a decrease of the optical anisotropy of the MT tip and thus increased presence of protofilaments in a curved conformation at the MT tip. The high temporal resolution of our imaging revealed a discrete nature of the MT disassembly and enabled the detection of individual tubulin oligomers detaching from the MT. The discrete nature of MT disassembly exposed periods in which no tubulin mass detached from the MT and the amount of isotropically-scattering tubulin structures at the MT tip was low. Strikingly, in the periods between individual unbinding events, we occasionally detected, in the absence of soluble tubulin, a momentary reversal of the apparent disassembly rate. The detected decrease of isotropic signal coinciding with these periods of stalled disassembly suggests that protofilaments of a disassembling MT, which are thought of as frayed, can transiently form a blunt and more closed structure. We hypothesize that these periods of stalled disassembly can act as moments enabling the switch from MT disassembly into MT assembly, termed rescue. We anticipate our experimental approach to become a vantage point for further experiments elucidating the functions of regulators of MT dynamics from a mechanistic point of view. In more general terms, we foresee the potential of our method for the investigation of anisotropy fluctuations on a single-protein level, providing new insight into the single-protein dynamics and super-resolution microscopy based solely on scattering changes of dynamic proteins.

## 4. Methods

### Reagents

Unless stated otherwise, all reagents were purchased from Sigma–Aldrich and used as received without further purification.

### Polymerization of double-stabilized MTs

Porcine brain tubulin was isolated according to a previously reported method.^[40]^ Microtubules were polymerized in the presence of GMPCPP, a slowly hydrolyzable analog of GTP, which protects the MTs from spontaneous depolymerization. Taxol was used in the MT solution to further enhance MT stability. First, a mixture containing 0.2 mg/mL tubulin, 1.0 mM GMPCPP (Jena Bioscience, Germany) and 1.0 mM MgCl_2_ in BRB80 (80 mM PIPES, 1 mM MgCl_2_, 1 mM EGTA, pH 6.9) was prepared and incubated at 37 °C for 2 hours. Then, the mixture was centrifuged at 12000 rpm for 20 minutes to separate the polymerized MTs from unbound dimers. The supernatant was removed and the pellet was resuspended in a BRB80 buffer containing 10 μM Taxol. These double-stabilized MTs were stored at room temperature until used in the experiments.

### Polymerization of GMPCPP-capped GDP MTs

The GMPCPP-capped MTs comprised a GDP body with both ends stabilized with GMPCPP caps to protect the GDP MTs from spontaneous depolymerization. First, the GDP MTs were grown by mixing 1.25 μL of a polymerization solution (BRB80, 25 % v/v of DMSO, 5 mM GTP, and 20 mM MgCl_2_) with 5 μL of a 4 mg/mL tubulin solution. This mixture was incubated at 37 °C for 2 h. After this, a 75 μL GMPCPP cap polymerization solution (BRB80, 1.33 mM GMPCPP, and 1.33 mM MgCl_2_) was heated to 37 °C, mixed with 5 μL of 4 mg/mL tubulin solution. Then, the GMPCPP-cap solution was added to the GDP MTs polymerization solution and the mixture was kept at 37 °C for 2 minutes. Finally, the GMPCPP-capped GDP MTs were stored at room temperature for no longer than 3 hours before they were used in the experiments.

### Flow channel assembly

All experiments involving MTs were performed in custom flow channels made of 22×22 mm (bottom, 1.5H thickness, Marienfeld, Germany) and 18×18 mm (top) coverslips separated by an ~100 μm thick parafilm spacer. First, both coverslips were rinsed with ethanol and ultrapure deionized water. After drying with N_2_, both coverslips were cleaned with oxygen plasma for 2 minutes. The bottom coverslip was silanized by dipping in acetone for 10 s and subsequent immersion in a 2 wt. % solution of 3-aminopropyl triethoxysilane (APTES) in acetone for another 10 s. Finally, the coverslip was rinsed with acetone and dried by a stream of N_2_. Next, two parallel stripes of parafilm were introduced between the top and bottom coverslips and heated on a hot plate to seal both sides of the channel.

### Kinesin-1 preparation

The DNA of the human kinesin-1 variant devoid of cysteine residue-encoding codons (Addgene #24430) consisted of amino acids 1 to 560 of KIF5B encoding a C-terminal 6xHis-tag^[41]^. A cysteine residue was re-introduced to amino acid 55 by point mutation from serine to cysteine performing site-directed mutagenesis. The mutated DNA was then expressed in *Escherichia coli* strain BL21(DE3) overnight at 18 °C after induction with 0.2 mM isopropyl-β-D-thiogalactopyranoside and in the presence of the respective antibiotic. The bacterial cells were harvested by centrifugation at 4000 g for 10 minutes at 4 °C, washed in PBS, and stored at −80 °C for further use. For cell lysis, the cell pellet was homogenized in lysis buffer (50 mM Na-phosphate buffer, pH 7.5, 5 % v/v glycerol, 300 mM KCl, 1 mM MgCl_2_, 0.1 % v/v Tween20, 10 mM BME, 0.1 mM ATP) supplemented with 30 mM imidazole, Protease Inhibitor Cocktail (cOmplete, EDTA free, Roche) and benzonase to a final concentration of 25 units/mL and lysed by a pulsed sonication for 5 minutes (20 s pulses with 24 W min^−1^). The cell lysis was further assisted by the addition of Igepal-630 to a final concentration of 0.2 % v/v. The cell lysate was separated from insoluble material by centrifugation at 40,000 g for 60 minutes at 4 °C and subsequently incubated with a lysis buffer-equilibrated Ni-NTA column (HisPur Ni-NTA Superflow Agarose, Pierce, VWR) for 2 h at 4 °C on a rotator for affinity chromatography via the 6xHis-tag. After washing the Ni-NTA column with wash buffer (His Trap buffer supplemented with 60 mM imidazole), the protein was eluted from the Ni-NTA column with an elution buffer (His Trap buffer supplemented with 300 mM imidazole). Fractions containing the protein of interest were pooled and concentrated using an Amicon ultracentrifuge filter and flash-frozen in liquid nitrogen.

### Functionalization of glass surface for the dynamic monitoring of MT disassembly

To immobilize the GMPCPP-capped GDP MTs on the surface of the flow channel, a 1 mg/mL β-casein solution in BRB80 was introduced into the channel for 15 minutes to adsorb on the APTES surface. Then, the unbound β-casein was removed with the attachment buffer (0.4 mM AMP-PNP and 10 mM DTT in BRB80). The next step comprised the injection of a 75 μg/mL kinesin solution (diluted in attachment buffer) for 5 minutes. The kinesin is known to bind casein surfaces, with the heads pointing at the solution.^[42]^ Then, 10 μL of the MT solution was injected into the channel and incubated for 5 minutes to anchor MTs to the kinesin heads in the presence of AMP-PNP (non-hydrolyzable ATP analog). Unbound MTs were removed by flushing the chamber with 20 μL of the attachment buffer.

### Functionalization of glass surface for the gliding of double-stabilized MTs

Experiments with double-stabilized MTs gliding on the surface were performed using kinesin-functionalized surfaces prepared following the same procedure as described above (monitoring of MT disassembly). Once the MTs were attached, 20 μL of buffer containing ATP (1.0 mM ATP and 10 mM DTT in BRB80) was injected into the channel to provide the energy for the kinesin-propelled μT movement along the surface.

### Functionalization of glass surface for the static measurement of MT scattering anisotropy

To immobilize the double-stabilized MTs, the flow channel with bottom APTES-coated coverslip was filled with 10 μL of the solution containing double-stabilized MTs in BRB80. After 1 minute, the unbound MTs were removed by injecting BRB80 into the channel.

### Preparation of dielectric nanowires

Dielectric nanowires were prepared using an electron beam lithography (EBL) system (eLINE Plus, Raith, Germany) on a glass coverslip. First, a thin layer of negative electron resist (XR-1541, Dow Corning, USA) was spin-coated on top of the cleaned glass surface and baked at 138 °C for 180 s using a hotplate. To enable charge dissipation during the e-beam exposure, a thin layer of conductive polymer (aquaSAVE, Mitsubishi Chemical, Japan) was spin-coated on top of the resist and baked at 138 °C for 60 s on a hotplate. A series of lines were patterned with a current dose optimized to yield nanowires with widths of approximately 20 nm. After the exposure, the conductive film was removed by rinsing in deionized water followed by the development of the pattern in 25% tetramethylammonium hydroxide (TMAH) developer for 35 s.

### Finite-difference time-domain model of anisotropic scattering

A commercial finite-difference time-domain method (FDTD) package (FDTD Solutions from Lumerical, Canada) was used for all calculations of scattering anisotropy. A scattering object, i.e., a model of MT or nanowire was positioned near the interface of the glass and water (MT) or glass and air (nanowire) and illuminated using a total-field scattered-field (TSFS) source (see Figure S5a). Refractive indices of materials used in the model were *n_water_* = 1.33, *n_glass_* = 1.51, *n_MT_* = 1.64^[43]^’ and *n_nanowire_* = 1.41. The power of the back-scattered field was read from a frequency-domain field and power (DFT) monitor placed outside the TSFS source in the direction opposite to the light propagation. The nanowire model had a rectangular crosssection of 80×25 nm and fine sub-mesh with 1×1×5 nm cell size was used. The MT was modeled as a hollow cylinder composed of 13 protofilaments of tubulins approximated as a chain of spheres with a 4 nm diameter and refractive index *n_MT_* (Figure S5). The outer diameter of the MT structure was 25 nm and a cubic mesh was used in the vicinity of the MT with a unit cell of 0.3 nm. Two simulations for both MT and a nanowire were run, each for the polarization parallel and perpendicular to the longest dimension of the scattering object. The ratio of the scattering amplitudes s^‖^/s^⊥^ was calculated as the square root of the ratio of the back-scattered powers.

With similar parameters of the FDTD simulation, the influence of the shape of a single protofilament on its scattering anisotropy was calculated (see Figure S5b).

### Monitoring the dynamics of MT disassembly with a polarization-sensitive iSCAT microscope

A simplified layout of the microscope used for polarization-sensitive iSCAT is shown in Figure 1a. A plane of polarization of the light emitted from a laser (λ = 660 nm, Cobolt Flamenco, Sweden) was rotated by 45° using a λ/2 waveplate to achieve even illumination in both polarization channels. Incident light was then focused (f = 500 mm) on the back focal plane (BFP) of the objective lens (NA = 1.3, UPLFLN100XOI2 from Olympus, Japan) resulting in a wide-field plane-wave illumination of the scattering sample contained in the flow channel. The same objective lens was used to collect the reflected (reference) and scattered waves. To achieve identical focus in the two polarization-channels and synchronous image acquisition, we used a configuration with a Wollaston prism, splitting the images in the two polarizations onto different areas of the same camera chip (MV1-D1024E, Photonfocus AG, Switzerland). The magnification of the microscope was 277×, corresponding to 38-nm-size of the image pixel.

A collimated beam from the second laser (*λ* = 532 nm, Gem 532, Laser Quantum, UK) was introduced into the common path of the microscope using a motorized retractable pellicle beam-splitter and used as the laser scissors. When the pellicle beam-splitter was in position, the green laser beam (100–200 mW) was focused on a part of an MT resulting in a photo-damage of an MT segment and initiating a disassembly of both freshly exposed ends. After exposing the MT to the laser scissors for 0.5 s, the pellicle beam-splitter was retracted and the MT disassembly was recorded for 8–16 seconds at the two orthogonal polarizations simultaneously in each camera frame (128 × 128 pixels) at up to 2900 frames per second.

When in focus, the MT contrast 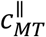 taken at the polarization parallel to the MT was higher than the contrast 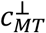 observed in the perpendicular polarization, which is consistent with the FDTD model (see Figure 2e). For an MT ideally aligned with one of the polarization planes of the microscope, the ratio 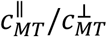 corresponds to the numerically calculated anisotropy parameter s^‖^/s^⊥^ (see Figure 2f). Due to the random angular misalignment of the MT orientation, the measured ratio 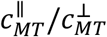 between 1.2 and 1.46 was slightly lower than the calculated value. However, this effect had a negligible impact on quantifying relative changes in the scattering anisotropy of a single MT.

### Two-polarization image alignment

The images from the two camera regions were overlapped with a sub-pixel resolution. In the alignment procedure, a set of incremental images was obtained by subtracting selected images of the same MT acquired with a 1 s delay. The incremental image selected for the alignment thus represents referenced images of small segments (approximately 220 nm long) of MT at ten different positions along the disassembling trajectory. The translation vector was then determined using the Lucas–Kanade method^[44]^ implemented in the image alignment toolbox in Matlab for each incremental image and the mean value was used for the alignment. The overall alignment error obtained from the set of calculated translation vectors was smaller than 5 nm (based on peak-to-peak variation).

### Construction of kymographs and accumulation of spatial profiles

Kymographs were constructed from longitudinal cross-sections of the MT image (linewidth of 3 pixels, i.e., ~110 nm). The averaged spatial profile of each MT end was retrieved from the particular kymograph corresponding to different timestamps in each polarization. A sigmoidal fit was used to identify centers of the profile slopes for each profile in one of the polarizations. Profiles taken at different time stamps were overlapped based on the offset of the fitted profile centers. To preserve the anisotropy-encoded information in the MT end profiles, the same offset was used for both polarizations. Finally, the average spatial profile (Figure 3d) was accumulated by overlapping the profiles from the individual experiments. Again, profiles in both polarizations were aligned based on offsets between the centers of the sigmoidal slopes fitted to all of the profiles in one selected polarization.

### Fitting the profile of disassembling MT scattering anisotropy

A simplified model of the nanoscopic scattering anisotropy profile of the disassembling MT was developed. A one-dimensional profile of the scattering cross-section along the MT was modeled as a product of the specific mass of the MT and its local polarizability α. The specific mass was considered constant along the whole length of the MT, whereas the polarizability was considered to be uniform in a shorter interval of the MT length, and the polarizability *α*^‖^ parallel to the MT was considered higher than the polarizability *α*^⊥^ perpendicular to the MT. A region of linear anisotropy loss characterized by a half-length of the linear gradient *d_depol_* was considered near the MT end, in which the local polarizability *α*^‖^ decreased linearly, while the *α*^⊥^ increased linearly towards the MT end at the same rate. This resulted in a linear decrease of the MT anisotropy from *α*^‖^ − *α*^⊥^ > 0 at a distance of 2*d_depol_* from the MT end towards *α*^‖^ − *α*^⊥^ = 0 at the MT end (see Figure S4d). The considered model was used to reconstruct the apparent far-field image profiles of MT in the parallel and perpendicular polarization. To suppress the effect of the PSF shape on the localization of the profile edge in the two detected polarizations, model profiles of scattering cross-sections in both polarizations were convolved with the combined PSF, obtained by convolving the two PSFs extracted from images measured in both polarizations. Finally, simulated mean and differential profiles of far-field contrast were calculated from the simulated profiles in both polarizations and fitted to the respective experimental profiles convolved with the PSF of the orthogonally polarized channel. The fitting was done in Python using a minimize package with the Nelder–Mead algorithm.^[45]^ The parameters *α*^‖^, *α*^⊥^ and *d_depol_* were considered as the free-fit parameters in the iterative algorithm. The effect of the free tubulin in solution of the measured scattering anisotropy near the MT tip was found to be negligible based on the high diffusion coefficient of tubulin.^[46]^ A more detailed discussion is provided in the Supporting Information.

An identical fitting algorithm was applied to the profile of scattering anisotropy accumulated from all disassembling MTs (Figure S4a,d) and two control structures, gliding double-stabilized MTs (Figure S4b,e), and dielectric nanowires (Figure S4c,f).

### Determination of the disassembly rate

The immediate velocities of the MT ends (both disassembling or gliding stable MT) were determined from a linear fit of the measured edge position within the 140 ms range. The localization precision of the MT edge was found to be *σ_x_* = 2.6 nm, which translates into a standard deviation of the calculated velocity of approximately *σ_vel_* = 9 nm/s.

### Analysis of the incremental images of the disassembling MT

Based on the width of the experimental PSF of *FWHM,_PSF_* = 350 nm, the number of tubulin dimers of single MT within a diffraction-limited volume is approximately *n_dimers_* = 570. The iSCAT contrast of the whole MT in our microscope (at λ = 660 nm) was *c_MT_* ≈ 3%. This results in a change of contrast of *Δc_dimer_* = *c_MT ndimers_^−1^* ≈ 0.005% when a single tubulin dimer unbinds from the MT.

The localization precision of the incremental peak is limited by the shot noise of the experiment, σ_loc,peak_ ~ N_photons_^−0.5^.^[47]^ We estimated the localization precision of the detected incremental peaks from the contrast and background noise to obtain the super-resolved image of the disassembled MT segment in Figure 6f,g. The incremental images were found to be shotnoise limited and any mechanical instabilities or drifts were much slower than the framerate.

### Detection of the mass displacement pattern in the incremental images

In each measured frame of disassembling MTs, we detected the position of the tip(s) of the MT based on the maximum spatial gradient of the contrast in the direction of the MT axis. We have calculated time incremental images with a 10-frames binning window (3.5 ms) meaning that for each timestamp we averaged the 10 preceding frames and subtracted it from the average of the 10 subsequent frames. We cropped the incremental images concerning the immediate position of the tip of the MT and rotated them in such a way that the direction of the MT disassembly was the same for all MT tips analyzed. We smoothed the incremental images with a Gaussian kernel of 2-pixel halfwidth. In the smoothed incremental images obtained in the polarization parallel to the MT axis, we detected all frames showing the mass displacement pattern, i.e., negative values at PSF halfwidth distance (4 pixels) from the MT end in the direction of the MT disassembly and positive values in the opposite direction and we used only the frames that had local maxima of the contrast pattern in the vicinity of 10 frames. Finally, we selected and averaged the frames measured in the perpendicular polarization based on the detection in the corresponding parallel polarization. The images in Figure 6 g-h were averaged from thousands of detected frames and the mean detection rate was approximately 180 per second for the disassembling MT (Fig. 6g) and 95 patterns per second for the gliding MT (Fig. 6h). This indicated that approximately 100 detections per second (all in the case of the gliding MT) are false fluctuation uncorrelated between the parallel and perpendicular polarization and indeed average out in Figure 6h. However, approximately 50% of the detected patterns (80 per second) in the case of the disassembling MT simultaneously appeared in both polarizations and could be detected and averaged even though they appeared considerably below the shot-noise limited background.

## Author Contributions

M.V. developed the optical setup, performed all the experiments and FDTD simulations; M.V, M.P. and Ł.B. processed the data; A.G.M. and K.H. developed the MT assay and laser scissors, V.H. generated the proteins, M.P., Z.L. and M.B. conceived the research; M.V., M.P., and Z.L. interpreted results and co-wrote the paper.

## Acknowledgments

This work was funded by the Ministry of Education, Youth and Sports of the Czech Republic under the project LL1602, the Czech Science Foundation under the project 18-19705S, the Introduction of New Research Methods to BIOCEV (CZ.1.05/2.1.00/19.0390) project from the ERDF, and the institutional support from the CAS (RVO: 86652036). The authors would like to thank David Palounek for his help with the experiments, and to Yulia Bobrova, Petr Dvořák, and Petra Lebrušková for technical support.

## Supporting Information

### The contrast in polarization-sensitive iSCAT microscope

Our dual-polarization iSCAT microscope measures the images of a scatterer projected into two orthogonal polarization planes (denoted as parallel, “‖” and perpendicular, “⊥”) of the microscope. The contrast in a standard iSCAT microscope is defined as:^[27]^ 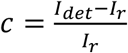, where *I_r_* is the intensity of the reference wave and the detected intensity 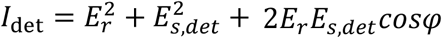. For small scatterers such as single MT, the pure scattering term 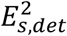 may be neglected. Maximum contrast then occurs when *cos φ* ≈ ±1 and the contrast is: 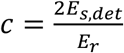 In our dual-polarization iSCAT microscope, the images are detected at two orthogonal polarizations denoted as ‖,_det_ and ⊥,_det_ (see Figure S1).

**Figure S1.**
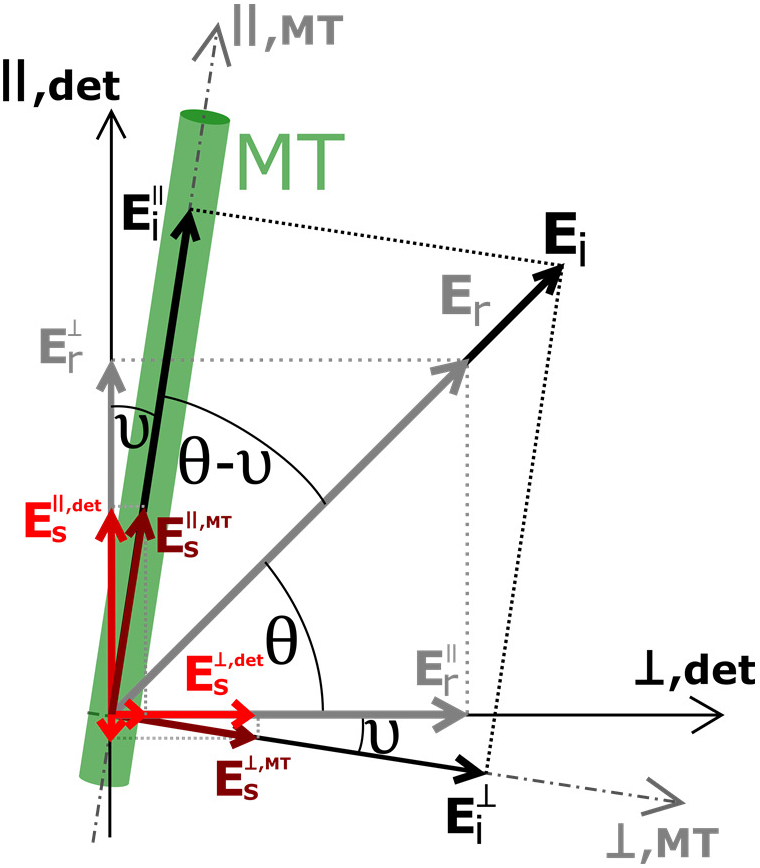
The geometry of the field scattered from MT and the reference field in the dual-polarization iSCAT microscope. Magnitudes of the incident (*E_i_*), reference (*E_r_*), and scattered (*E_s_*) fields are not to scale.

The incident wave was polarized linearly at 45 deg to both polarization directions ‖,_det_ and ⊥,_det_, i.e. *θ* = 45 deg to achieve even illumination on both polarization channels. In general, the MT is not perfectly parallel with ‖,_det_ axis, instead, it is misaligned by the angle υ. Next, we assume the MT is a uniaxial scatterer with distinct scattering amplitudes s^‖^ and s^⊥^ for the light polarized parallel and perpendicular to the MT, respectively. In such a case, the field scattered by the MT can be written as:

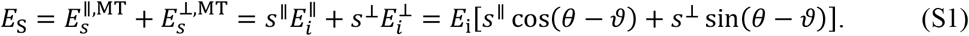

When the MT is aligned with ‖,_det_ (υ = 0), the measured contrasts c^‖^ and c^⊥^ are directly proportional to the MT scattering amplitudes s^‖^ and s^⊥^ as 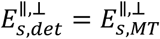. When υ ≠ 0, The detected fields are:

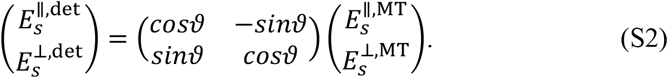

To keep the mixing of the 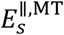 and 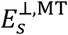 signals reasonably low, we have taken into account only MTs close to parallel to the ‖,_det_, i.e. υ ⪅ 10 deg. In the limit case of υ = 10, the detected anisotropy ratio 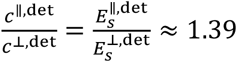, when the 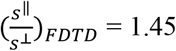 obtained from the numerical FDTD model (see section FDTD model of anisotropic scattering below) was plugged into the calculation (Equations S1 and S2). This means that even in the limit case of υ = 10 deg, the effect of the misalignment on the measured anisotropy ratio 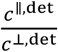 is that the detected ratio is only about 5% smaller than the real scattering anisotropy ratio of the MT, 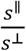.

### Characterization of the prepared nanowires

Nanowires fabricated by electron beam lithography (EBL) from HSQ were characterized using scanning electron microscopy (SEM) and atomic force microscopy (AFM) to measure their width and height, respectively. While the determination of height from AFM data is straightforward, nanowire widths measured from SEM images are affected by the gold film (25 nm thickness) deposited on top of the nanowires in order to facilitate the electron microscopy. The mean width of the nanowire measured from the SEM image was 35 nm. Considering the typical sidewalls angle of the evaporated layer of 12 deg we estimate the real width of a nanowire to be around 25 nm. Combined, based on this characterization, we assume the prepared nanowires have a rectangular cross-section of 80 nm × 25 nm (height × width).

**Figure S2.**
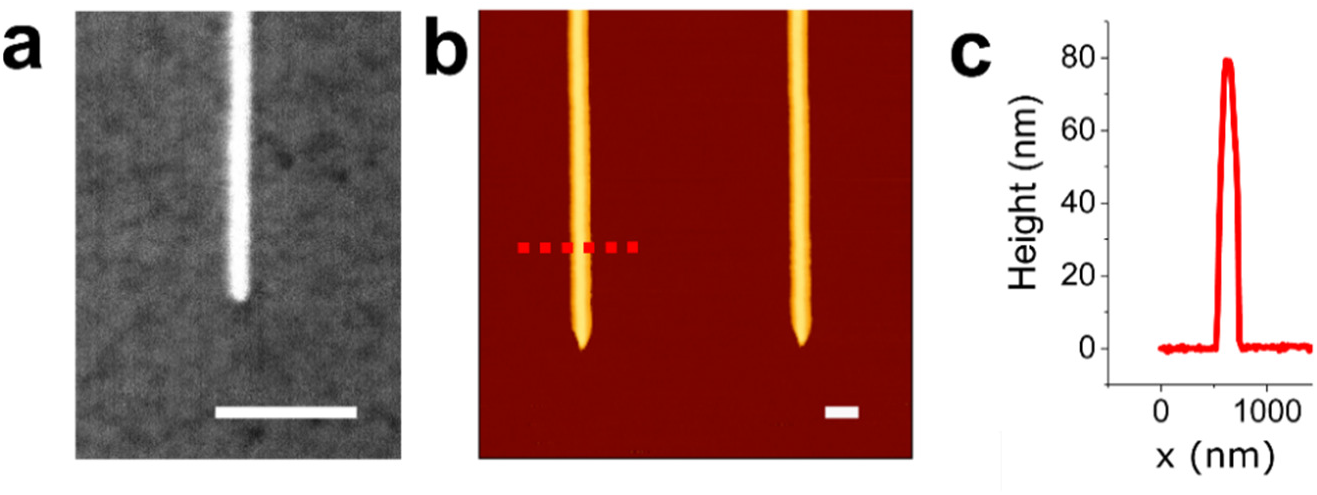
Characterization of the prepared nanowires. (**a**) SEM micrograph of a dielectric nanowire coated with 25 nm of gold. (**b**) AFM scan of two nanowires with the height crosssection. The length of scale bars corresponds to 300 nm.

### Linearized model of the anisotropy profiles of individual disassembling MT ends

Time series of iSCAT images were acquired in two orthogonal polarizations and the corresponding profiles of the MT contrast along the MT near each MT tip (N=16 MT tips were processed) were obtained using a procedure described in Methods. All time-averaged spatial profiles of the mean and differential contrast from individual MT kymographs are presented in Figure S3. An example of fitted curves (dash-dotted lines), based on a simple model of linear decrease of the anisotropy near the MT tip (see Figure S4d) are shown overlapped with the experimental profiles. The fitting procedure is described in Methods.

**Figure S3.**
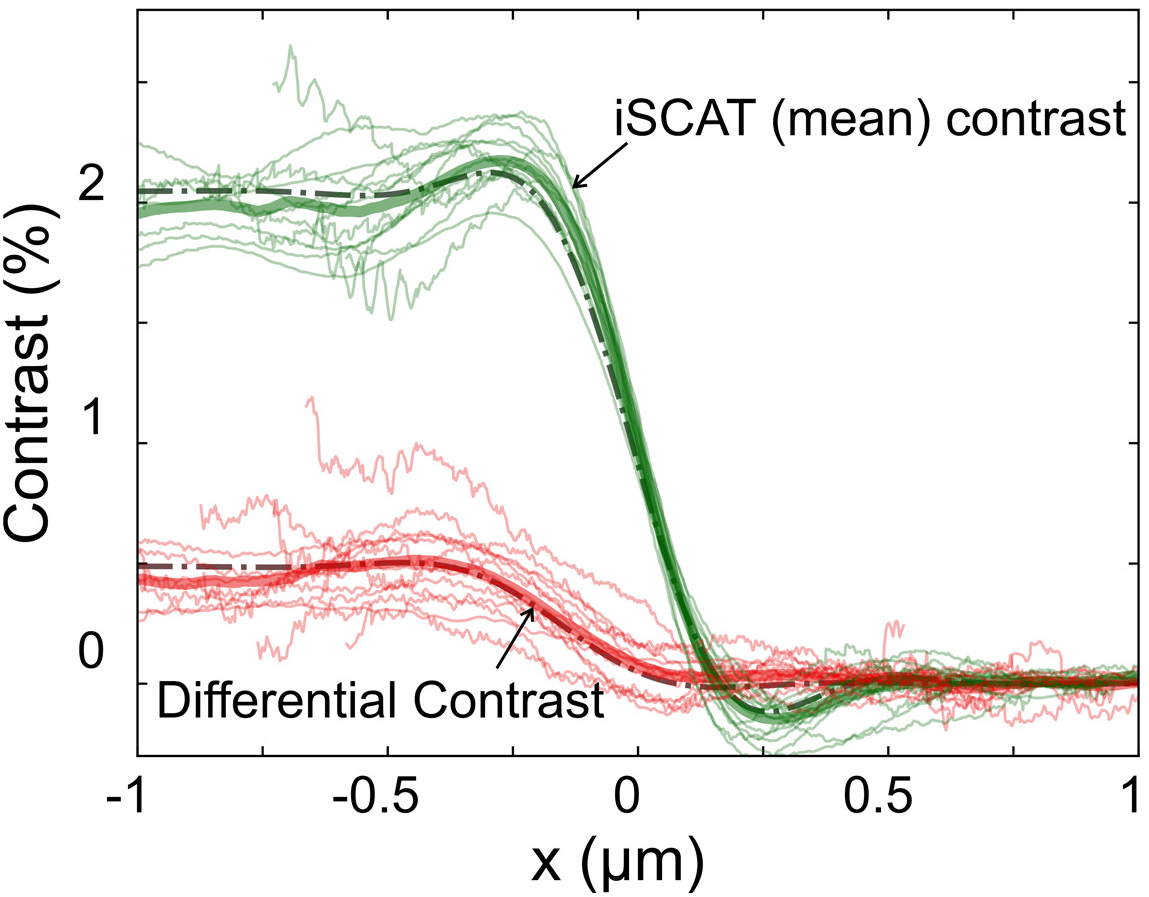
Spatial profiles of the mean (green lines) and differential (red lines) contrast measured from 14 disassembling MT ends. Profiles averaged from all 14 ends (thick solid lines) and fitted curves (dash-dotted lines) obtained from a simple model of MT mass and polarizability distribution are shown for comparison.

When averaged spatial profiles of mean and differential MT contrast is fitted with the profiles constructed as a convolution of the profile of polarizability along the MT convoluted with the microscope PSF, we obtain the parameter *d_depol_* as a free parameter of the fit. Here, we have obtained *d_depol,dMT_* = (90 ± 1) nm for the disassembling MT (Figure S4a,d), *d_depol,sMT_* = (25 ± 1) nm for stable MT (Figure S4b,e), and *d_depol,wire_* = (3 ± 1) nm (Figure S4c,f). Standard deviations in all cases were calculated from a randomized fit to fit variation. The values of *d_depol_* are in good agreement with average values of *Δx_c_* for all three studied structures (see Figure 3g) and provide a simple model of the anisotropy loss near the MT or nanowire ends.

**Figure S4.**
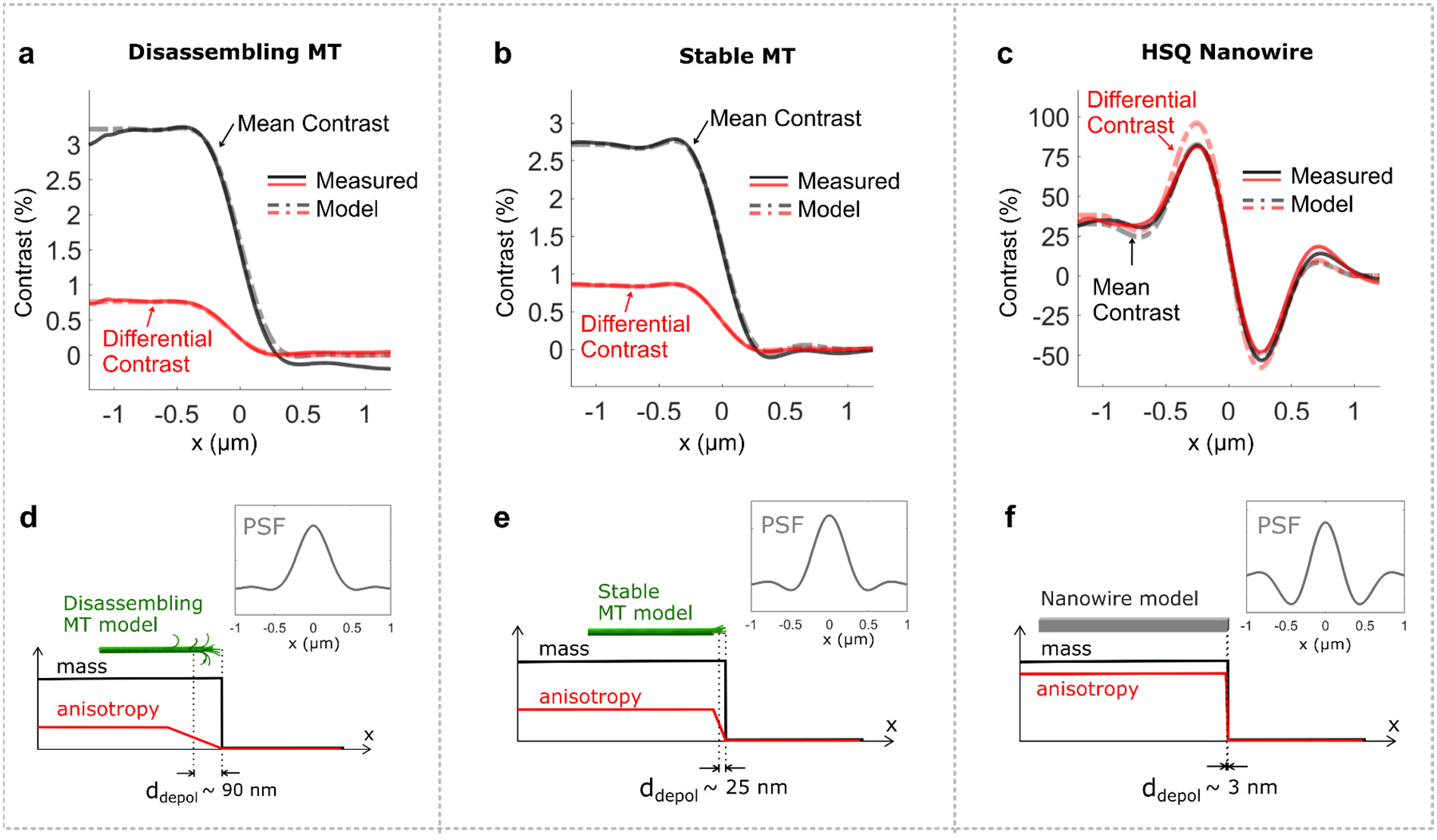
Model of the spatial profile of the anisotropy loss at the disassembling MT tip and control structures. Spatial profiles of the mean (black line) and differential (red line) iSCAT contrast averaged from (a) disassembling MT ends (N=16), (b) stable ends of gliding MTs (N=42), and (c) fabricated HSQ nanowires (N=20). A simple model of linearly decreasing anisotropy for (c) disassembling MT, (d) stable MT, and (e) nanowire and the PSFs extracted from the experimental data for each structure. Fitted profiles obtained using models in (d-f) are shown in (a-c) as dash-dotted lines.

### FDTD model of anisotropic scattering

An FDTD model was created to simulate the scattering anisotropy of the whole MT (Figure S5a). In this simulation, the tubulin molecules were modeled as isotropic spheres with a refractive index of 1.64.^[43]^ The resulting simulated scattering anisotropy was 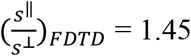.

We also performed a series of FDTD simulations comparing the scattering anisotropy of straight protofilament segment and circularly coiled protofilament segment of the same length at two different orientations to the illumination wave as illustrated in Figure S4b as cases (i), (ii), and (iii). The modeled protofilament segment comprised 12 spheres (4 nm in diameter) representing the tubulin molecules in all three modeled cases. The simulated cases correspond to (i) straight protofilament, (ii) protofilament curled “upwards”, i.e. along the direction of the microscope axis, and (iii) protofilament curled “sideways” from the MT shaft, i.e. in the plane perpendicular to the microscope axis. In each of these cases, the power of the back-scattered field detected by the frequency-domain field and power (DFT) monitor (see Methods) was acquired for the two polarizations of the TFSF (total-field scattered-field) source, parallel and perpendicular to the MT shaft (*x*-axis).). As a result, the respective ratios of the scattering amplitudes were: (i) (s^‖^/s^⊥^)_straight_ = 1.12 for straight protofilament, (ii) (s^‖^/s^⊥^)_upcurved_ = 1.05 for the “upwards” curved protofilament and (iii) (s^‖^/s^⊥^)_sidecurved_ = 1.00 for the “sideways” curved protofilament. These results clearly indicate that local anisotropy of the MT is affected by the shape of the protofilaments and the formation of coiled protofilaments or “ram’s horns” reduces the scattering anisotropy by 60 to 100 percent depending on their orientation to the probing light.

**Figure S5.**
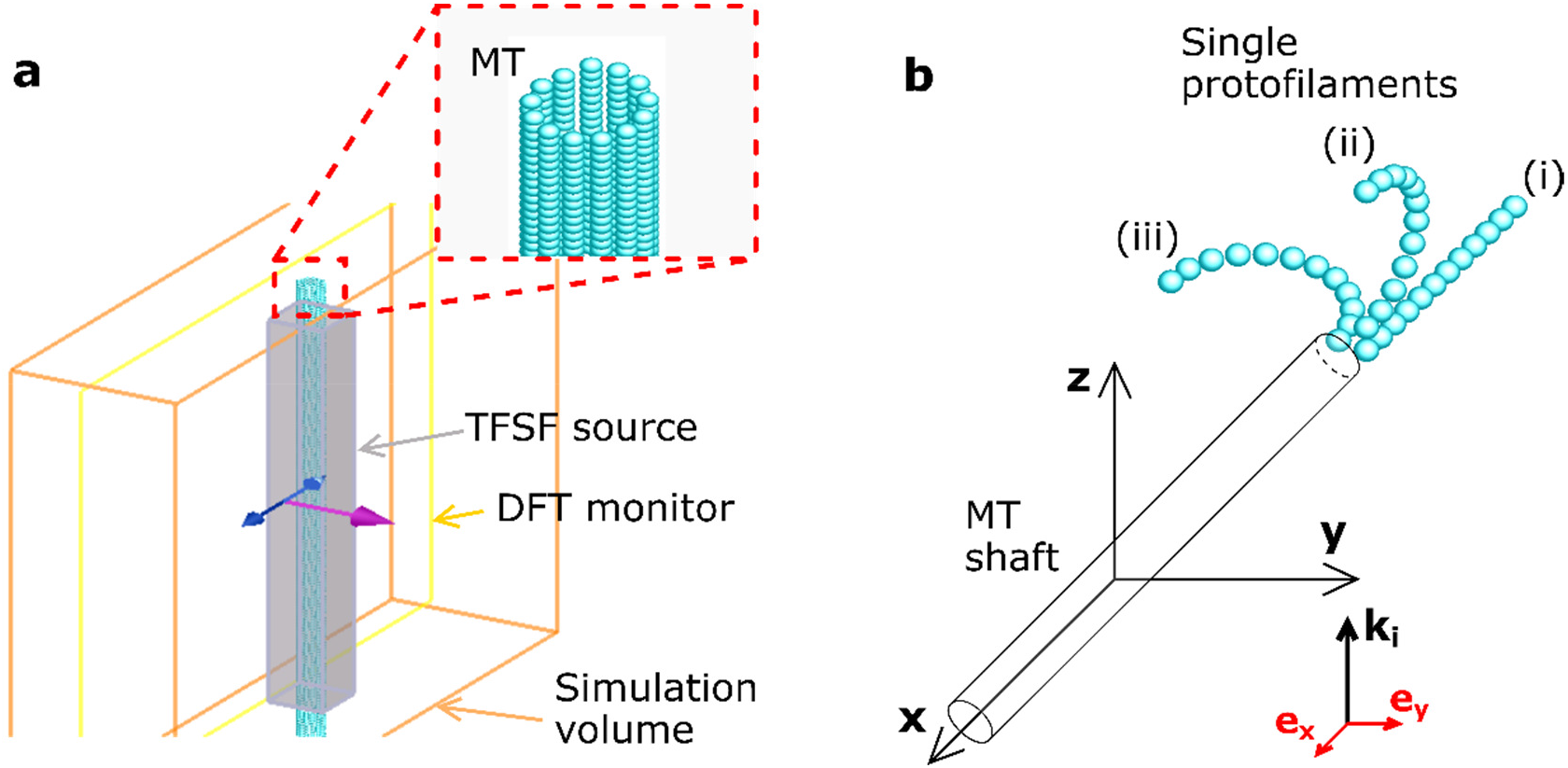
FDTD model of the MT. (**a**) The geometry of the FDTD simulation used in the calculation of the scattering anisotropy of stable MT. (**b**) The geometry of single protofilament segments used in FDTD simulations. (i) straight protofilament; (ii) protofilament curled into the plane parallel with the direction of the incident light beam (indicated by ***k_i_*** ‖ ***z*)** and (iii) protofilament curled into the plane perpendicular to the direction of the incident light beam.

### Effect of the free tubulin on the measured scattering anisotropy

Apart from the structural changes in the shape of protofilament segments, another possible cause of an apparent anisotropy loss may originate in the presence of free tubulin dimers or protofilament segments released from the disassembling MT end contributing to the overall scattering signal. When released, tubulin diffuses freely in solution with a diffusion coefficient of D = 4.3 × 10^−11^m^2^ s^−1^.^[46]^ Considering the diffusion in three dimensions, the mean square displacement of a tubulin dimer in time *t* after the release will be 〈*r*^2^〉 = 6*Dt*. Therefore, the mean time required for a single tubulin dimer to escape from a spherical volume of 1 μm in diameter is t ≈ 1 ms. Considering the average disassembly rate of 0.5 tubulin dimers per millisecond, the average contribution to the decrease in the apparent anisotropy due to this effect would count as one out of more than 1600 tubulin dimers present in the volume of a 1 μm in diameter. As the experimental effect of the anisotropy change reads more than 30%, we consider the effect of the recently unbound tubulin dimer negligible.

### An estimate of the frequency of long ram’s horns unbinding

If we assume an average apparent speed of disassembly of *v_disassembly_* =220nm s^−1^, the average rate at which the tubulin dimers leaves the MT shaft is *V_dimers_* ≈ 360 s^−1^ (*v_dimers_* = *v_disassembly nPFldimer_^−1^*, where *n_PF_* = 13 is the number of protofilaments in MT and *l_dimer_* = 8 nm is a length of one tubulin dimer). Let us assume that tubulin dimers can unbind from curled protofilament (PF) (ram’s horns) only, i.e. when lateral bonds between PFs are already broken. The simultaneous break of both lateral and longitudinal bonds is much less probable. In the case of an average length of ram’s horn of ~100 nm (full circle of 30 nm in diameter), there are approximately 12 tubulin dimers per ram’s horn. Next, we assume the probability of breaking the longitudinal bond, i.e. between tubulin dimers in one PF, to be equal for all longitudinal bonds in that ram’s horn. This means any number of between 1 and 12 of tubulin dimers will leave the MT with the same probability. This means that the rate at which the segments with lengths between 3 and 12 dimers unbind the MT, is *v_segments,3-12_* = 41 s^−1^. This estimation does not take into account the non-linear distribution of the lengths of the formed ram’s horns as well as other factors. Surprisingly, the number of detected unbinding events within a *t_exp_* = 4.8-second-long experiment (Figure 6h) was 199, which is in line with the number of ≈ 197 segments (*v_segments,3-12_* × *t_exp_* s) obtained using the above-described assumptions.

### Statistics of the disassembly rates

Traces and immediate velocities of the one pair of the disassembling ends are shown in Figure S6a-c. Histograms of velocities for all measured disassembling MT ends are then summarized in Figure S6d. Vertical dashed lines separate pairs of plus and minus ends that arise after cutting the single MT with the laser scissors. Due to the random orientation of the MTs on the surface and similar disassembly rates of plus and minus ends,^[8]^ we could not undoubtedly assign the plus and minus polarity to the ends in the pairs. However, even without this information, we can see that the disassembly rates vary substantially between zero (or small negative values) and several hundred nm per second in the vast majority of experiments and both MT polarities.

**Figure S6.**
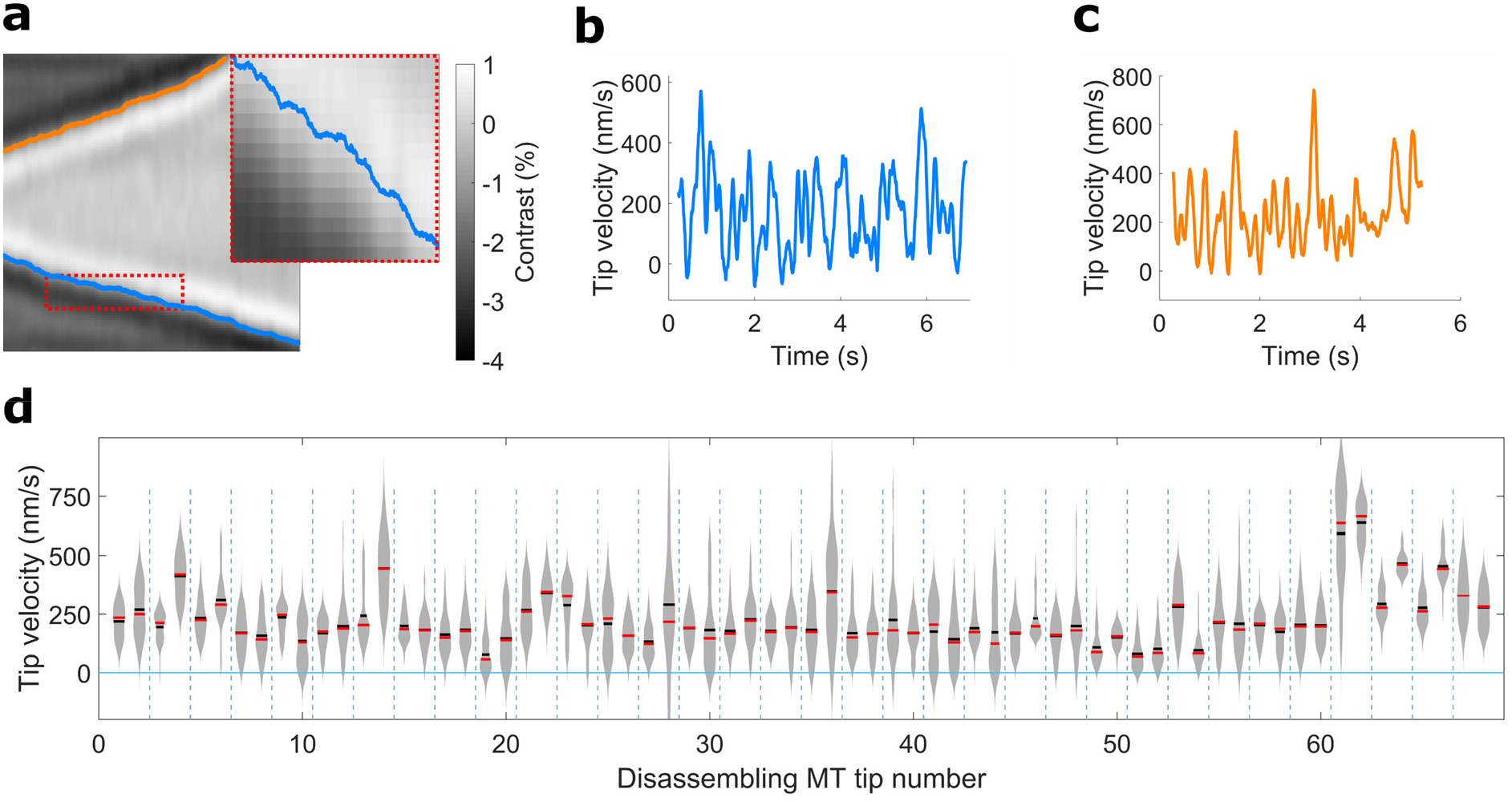
(**a**) Kymograph of the mean iSCAT signal with tracked positions of two receding ends of the disassembling MT. (**b**,**c**) immediate velocities of the receding ends calculated from positions of the ends in (**a**). (**d**) Histograms of the velocities of n=68 individual disassembling MT ends. Vertical lines separate individual experiments grouping the plus and the minus end of the same MT into a pair.

### SNR of the unbinding events of tubulin oligomers

As the iSCAT contrast of single tubulin dimer is only about 5 × 10^−5^, the detection of the events where tubulin unbinds from the MT tip is limited by the experimental noise. In our experiment, up to 6.6 × 10^8^ photons are detected per PSF, when 200 ke^−^ well depth of the camera pixel, *FWHM_PSF_* = 350 nm, 38 nm effective pixel size, and frame averaging of *N_bin_* = 50 are considered. This translates into shot-noise in the order of *σ_noise_* ≈ 4 × 10^−5^.

